# Investigating the role of root exudates in recruiting *Streptomyces* bacteria to the *Arabidopsis thaliana* root microbiome

**DOI:** 10.1101/2020.09.09.290742

**Authors:** Sarah F. Worsley, Michael Macey, Sam Prudence, Barrie Wilkinson, J. Colin Murrell, Matthew I. Hutchings

## Abstract

*Streptomyces* species are saprophytic soil bacteria that produce a diverse array of specialised metabolites, including half of all known antibiotics. They are also rhizobacteria and plant endophytes that can promote plant growth and protect against disease. Several studies have shown that streptomycetes are enriched in the rhizosphere and endosphere of the model plant *Arabidopsis thaliana*. Here, we set out to test the hypothesis that they are attracted to plant roots by root exudates, and specifically by the plant phytohormone salicylate, which they might use as a nutrient source. We confirmed a previously published report that salicylate over-producing *cpr5* plants are colonised more readily by streptomycetes but found that salicylate-deficient *sid2-2* and *pad4* plants had the same levels of root colonisation by *Streptomyces* bacteria as the wild-type plants. We then tested eight genome sequenced *Streptomyces* endophyte strains *in vitro* and found that none were attracted to or could grow on salicylate as a sole carbon source. We next used ^13^CO_2_ DNA stable isotope probing to test whether *Streptomyces* species can feed off a wider range of plant metabolites but found that *Streptomyces* bacteria were outcompeted by faster growing proteobacteria and did not incorporate photosynthetically fixed carbon into their DNA. We conclude that, given their saprotrophic nature and under conditions of high competition, streptomycetes most likely feed on more complex organic material shed by growing plant roots. Understanding the factors that impact the competitiveness of strains in the plant root microbiome could have consequences for the effective application of biocontrol strains.

**Importance:** *Streptomyces* bacteria are ubiquitous in soil but their role as rhizobacteria is less well studied. Our recent work demonstrated that streptomycetes isolated from *A. thaliana* roots can promote growth and protect against disease across plant species and that plant growth hormones can modulate the production of bioactive specialised metabolites by these strains. Here we used ^13^CO_2_ DNA stable isotope probing to identify which bacteria feed on plant metabolites in the *A. thaliana* rhizosphere and, for the first time, in the endosphere. We found that *Streptomyces* species are outcompeted for these metabolites by faster growing proteobacteria and instead likely subsist on more complex organic material such as cellulose derived from plant cell material that is shed from the roots. This work thus reveals the “winners and losers” in the battle between soil bacteria for plant metabolites and could inform the development of methods to apply streptomycetes as plant growth-promoting agents.

## Introduction

*Streptomyces* species are saprophytic soil bacteria that play an important ecological role as composters in soil and are prolific producers of antimicrobial compounds (1,2). The genus first evolved ca. 440 million years ago when plants began to colonise land and it has been suggested that their filamentous growth and diverse secondary metabolism may have evolved to enhance their ability to colonise plant roots (3). Several studies have reported that streptomycetes are abundant inside the roots of the model plant *Arabidopsis thaliana* (4–10) where they can have beneficial effects on growth (11) while others have shown that streptomycetes can protect crop plants such as strawberry, lettuce, rice and wheat against biotic and abiotic stressors, including drought, salt stress, and pathogenic infection (12,13). However, different plant genotypes are associated with distinctive root-associated microbial communities and not all plant species enrich streptomycetes in their roots, a notable example being barley (14). This suggests that either specific selection mechanisms exist that enable different plant hosts to recruit particular microbial species from the soil and / or that bacterial taxa such as *Streptomyces* species colonise some plants better than others.

In our previous work, we reported that three *Streptomyces* strains isolated from the roots of *A. thaliana* plants can be re-introduced to new plants to promote their growth both *in vitro* and in soil. Furthermore, one of these strains exhibited broad spectrum antifungal activity and, when its spores were used as a seed coating, it colonised the roots of bread wheat plants and protected them against the commercially important pathogen *Gaeumannomyces tritici*, the causative agent of take-all disease (11). Here we set out to test the hypothesis that streptomycetes are attracted to, and metabolise, components of the *A. thaliana* root exudate. Plants are known to release up to 40% of their photosynthetically fixed carbon into the surrounding soil via their roots and bacterial species in the soil have been shown to be attracted to, and be capable of metabolising, specific resources contained within these root exudates (15–17). In turn, it is thought that the plant host might establish a beneficial root microbiome by producing particular types of root exudate that attract microbial species with desirable metabolic traits (18,19). Recent studies have further suggested that defence phytohormones, which are accumulated by plants in response to pathogenic attack, may play a key role in modulating the establishment of the normal *A. thaliana* root microbiome, since mutant plants that are disrupted in these pathways accumulate significantly different microbial communities (9,10). In particular, the abundance of bacteria in the *A. thaliana* rhizosphere and roots has been shown to correlate with concentrations of the defence phytohormone salicylic acid (10). Plant growth hormones such as indole-3-acetic acid have also been shown to modulate the production of *Streptomyces* specialised metabolites which may aid their competitive establishment in the root microbiome (11).

Here we set out to test the hypothesis that streptomycetes are attracted by, and feed on, salicylate as well as other components of *A. thaliana* root exudates. We used genome sequenced *Streptomyces* endophyte strains that we isolated in a previous study (11), including three which have been shown to promote *A. thaliana* growth *in vitro* and in compost, and assessed their ability to colonise mutant *A. thaliana* plants affected in salicylate production. Our results indicate that salicylate does not attract or feed these *Streptomyces* species, at least under the conditions used in our experiments. Furthermore, ^13^CO_2_ DNA stable isotope probing (DNA-SIP) of wild-type *A. thaliana* plants grown in compost show that *Streptomyces* bacteria do not feed on root exudates and are instead outcompeted by faster growing proteobacteria. We propose that streptomycetes are more likely to feed on complex organic matter that is not labelled in the ^13^CO_2_ DNA-SIP experiments. This work is an important step in understanding the role of these bacteria in the plant rhizosphere and also reveals which members of the *A. thaliana* microbiome feed on plant metabolites.

## Results

### Root colonisation by *Streptomyces* bacteria is affected by plant genotype

A previous study reported that *A. thaliana cpr5* plants that constitutively produce salicylate have an altered root microbiota compared to wild-type plants, and that streptomycetes were better able to colonize these plants *in vitro*, due to their ability to metabolize salicylate (10). This suggests that salicylate might be directly responsible for recruiting these bacteria to the plant roots. To test this hypothesis, we compared root colonization efficiencies by *S. coelicolor* M145 and the root-associated strain *Streptomyces* M3 in wild-type *A. thaliana* Col-0 plants, *cpr5* plants as well as two other genotypes, *pad4* and *sid2-2* plants, that are deficient in salicylate production (Table 1) (20–23). To measure plant root colonization efficiency, we established root infection assays in which pre-germinated *Streptomyces* spores were used to coat *A. thaliana* seeds and were also added to the surrounding soil. *Streptomyces* strains M3 and *S. coelicolor* M145 (Table 1), were used because they have previously been observed to interact extensively with the *A. thaliana* roots using confocal microscopy (11). Both strains have been engineered to carry the apramycin resistance (*aac*) gene, enabling the selective re-isolation of these strains from the roots of soil grown plants and their subsequent enumeration on agar plates containing apramycin. Colonization was measured as colony forming units (cfu) retrieved per gram of plant root tissue. This method has been used previously to assess colonization efficiency by other streptomycete strains in the plant root microbiome (24,25). The results of these assays show that root colonization was significantly affected by plant genotype (Fig. 1), irrespective of the *Streptomyces* strain used as an inoculum; plant genotype had a significant effect on the log-transformed cfu g^−1^ (F_(3,39)_ = 6.17, *P* <0.01), whereas the strain-genotype interaction term was insignificant (F_(3,39)_ = 0.51, *P* = 0.68) in an ANOVA test. Indeed, consistent with previous findings (10), colonization by both strains significantly increased in the *cpr5* mutant plants which constitutively make salicylate (Fig. 1), compared to wild-type plants and the salicylate-deficient plants, *pad4* and *sid2-2 (P* < 0.05 in all Tukey’s HSD tests between *cpr5* and the other plant genotypes). However, we observed no significant difference in root colonization by M145 or M3 in *pad4* or *sid2-2* plants, compared to wild-type *A. thaliana* plants (*P* > 0.05 in Tukeys HSD tests), suggesting that reduced quantities of salicylate do not in turn correspond to a reduction in colonization by streptomycetes (Fig. 1). We note that the *cpr5* gene has a complex role in regulating pathways involved in plant growth, immunity, and senescence (21) and that the *cpr5* plants grown in our experiments were compromised in their growth compared to all the other plant genotypes tested here (Fig. 1). The altered development of *cpr5* plants has been observed in numerous other studies (21,26–29) and gives rise to the possibility that the observed increase in colonization by *Streptomyces* strains M145 and M3 was linked to other aspects of the complex phenotype of the *cpr5* plants and was not necessarily due to the higher levels of salicylate produced by these plants.

**Table 1.**
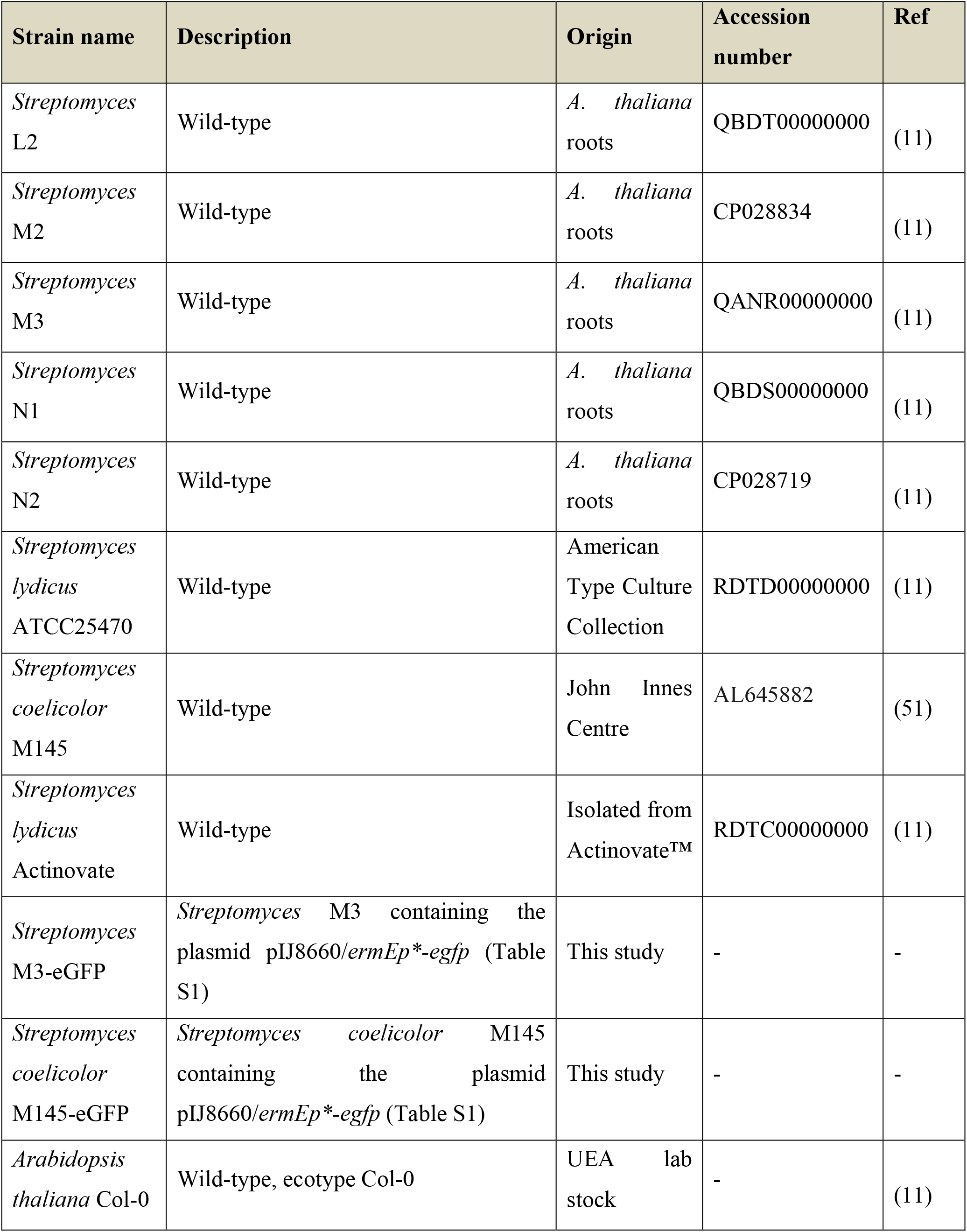
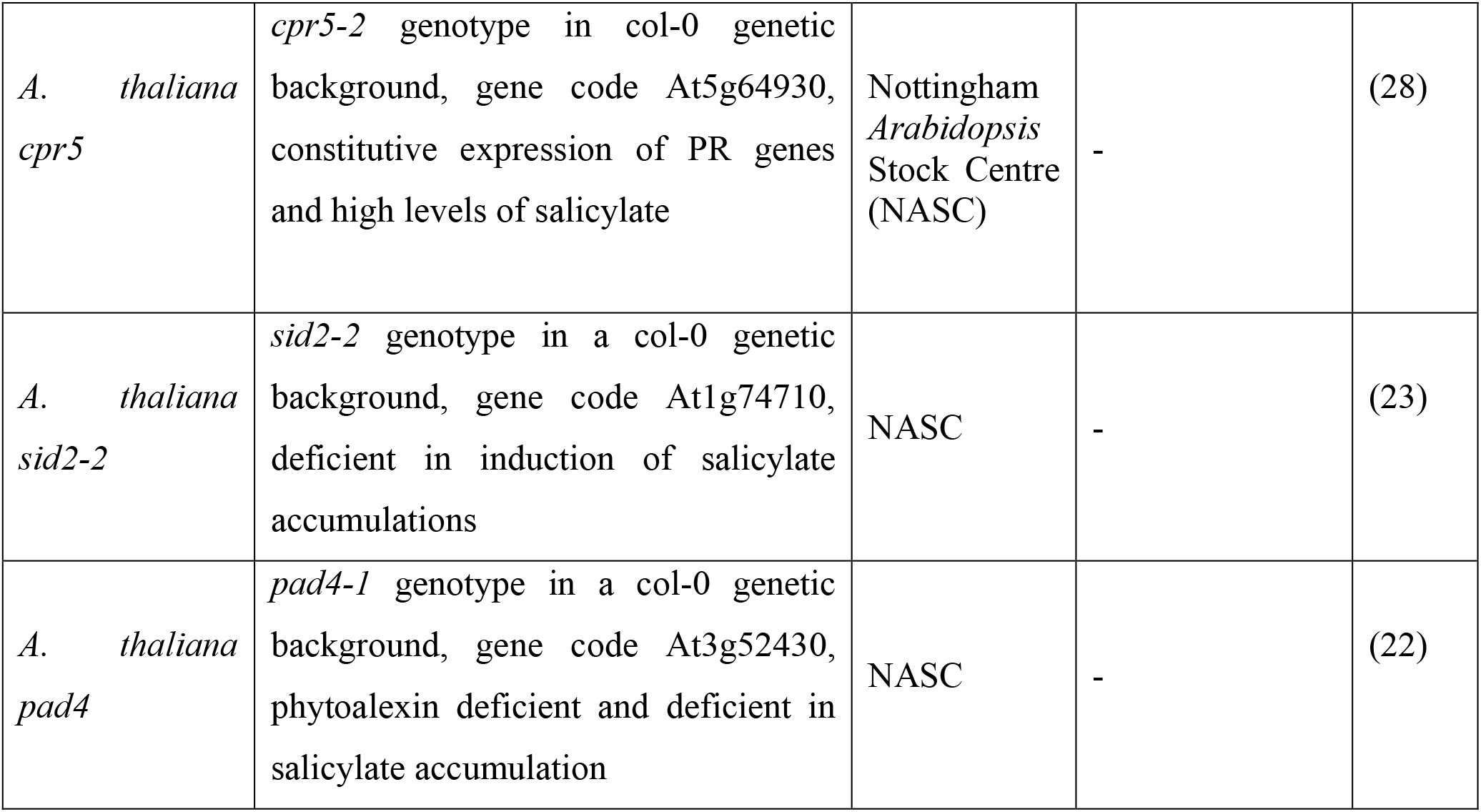
Bacterial strains and plants used in experiments.

**Figure 1.**
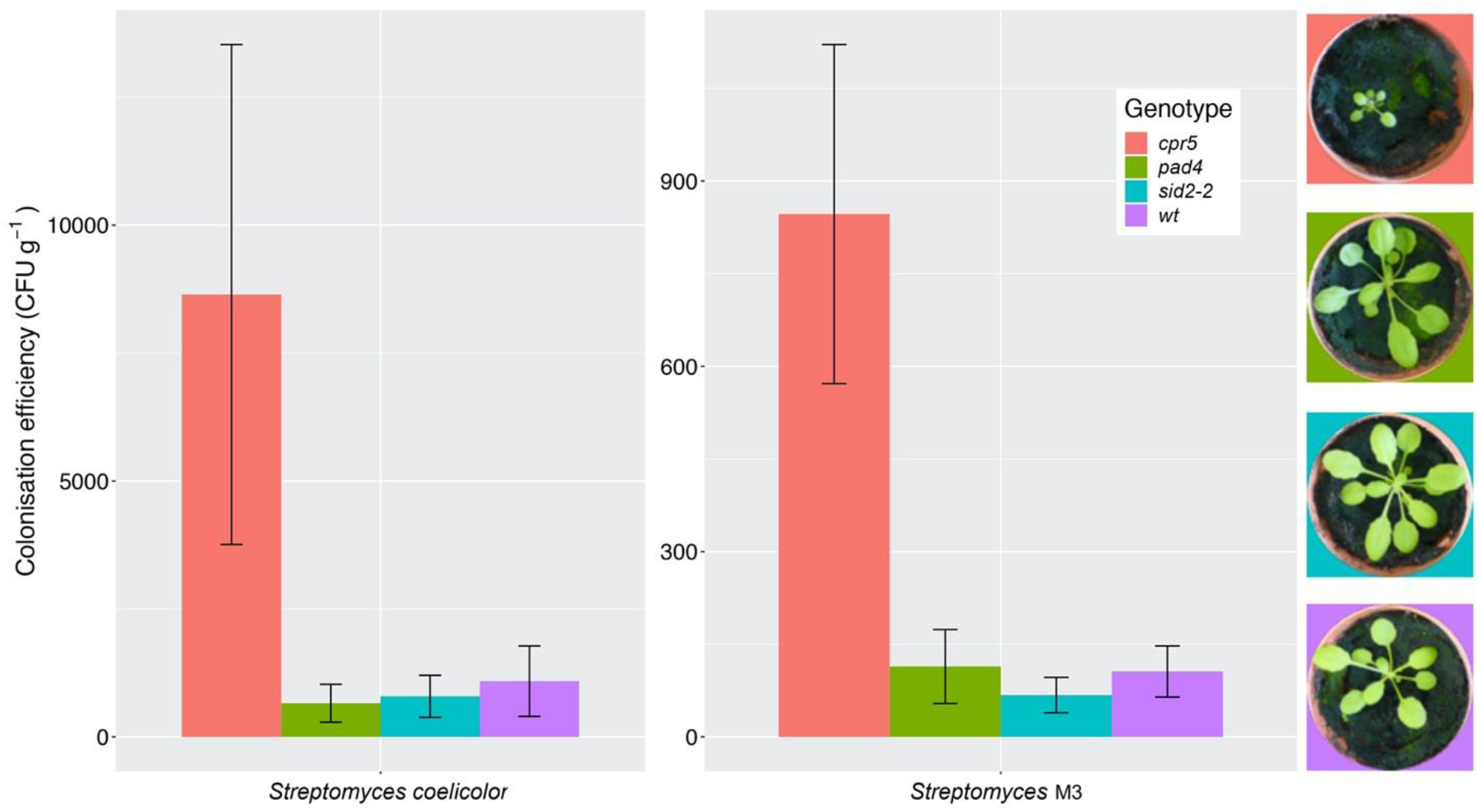
*Streptomyces coelicolor* M145 and the *A. thaliana* root endophyte strain *Streptomyces* M3 show increased root colonization of *Arabidopsis thaliana cpr5* plants, which constitutively produce salicylic acid, compared to *pad4* and *sid2-2* plants, which are deficient in salicylic acid production, and wild-type (wt) Col-0 plants. Root colonisation was measured as the average colony forming units (±SE) of *Streptomyces* bacteria per gram of root, that could be re-isolated from plants four weeks after germination. N = 6 plants per treatment. Plant growth phenotypes are shown for comparison.

### *Streptomyces* species are not attracted by and do not feed on salicylic acid *in vitro*

To test whether the streptomycete strains used in recolonization experiments can utilise salicylate as a sole-carbon source, we grew each strain on minimal agarose medium (MM) containing either sodium citrate, or their preferred carbon sources, mannitol, maltose or sucrose. We also tested seven additional strains, four of which were also isolated from *A. thaliana* roots, as well as three strains of *Streptomyces lydicus* which are known colonizers of plant roots and were genome-sequenced in a previous study (11). All the strains grew well on their preferred carbon source but none of the strains grew on MM plates containing salicylate as the sole carbon source (Fig. 2). This included *Streptomyces* M3 which had demonstrated increased levels of colonization in *A. thaliana cpr5* plants (Fig. 1). Thus, salicylate is not used as a carbon source by any of the *Streptomyces* strains used in our study, and this is supported by the absence of known SA degradation genes in the genomes of all these isolates (Table S1). Rather than acting as a nutrient source, some root exudate molecules may recruit bacteria to the root niche by acting as chemoattractants (30). In order to test whether salicylate is a chemoattractant for *Streptomyces* species, we grew our test strains on agar plates next to paper disks soaked in either 0.5 mM or 1 mM salicylate, or 0.1% (v/v) DMSO (solvent control) but observed no growth towards salicylate after 10 days (Fig. S1).

**Figure 2.**
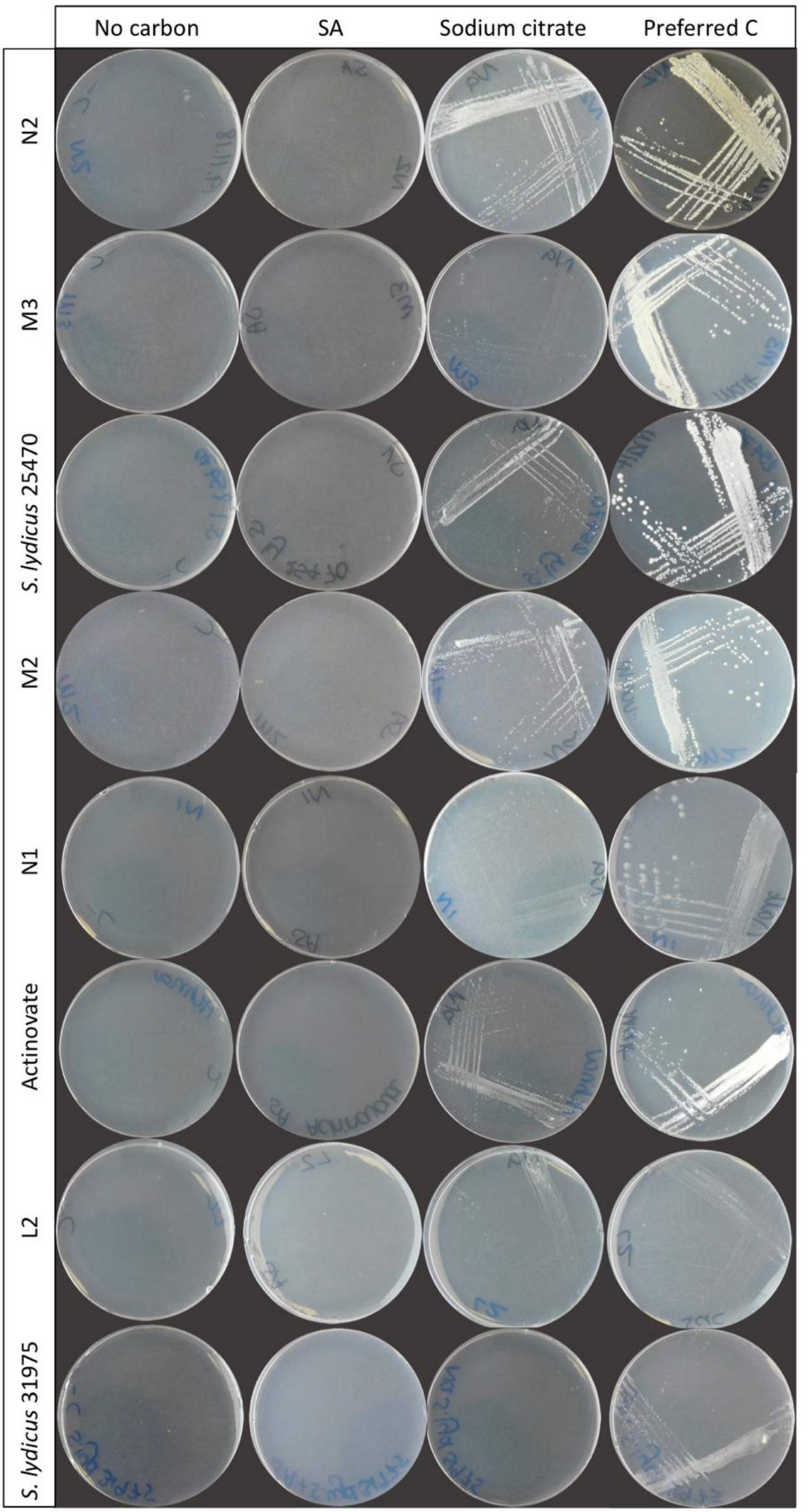
*Streptomyces* isolates (as indicated) grown on minimal medium agarose plates containing either no carbon, 0.5 mM salicylic acid (SA), 3.875 mM sodium citrate or their preferred carbon source.

To test whether altered salicylate levels might indirectly benefit streptomycetes by negatively modulating the levels of the other rhizobacteria in the soil and removing competition, we established soil microcosms in deep 12-well plates, which were wetted with either sterile water or 0.5 mM salicylate. Microcosms were inoculated with spores of *S. coelicolor* M145-eGFP or *Streptomyces* M3-eGFP, both of which had colonised *cpr5* plants more effectively than wild-type plants. Nine replicates of each treatment were run in parallel for each strain and the uninoculated control. Strains were recovered on apramycin-containing selective agar medium after 10 days. No apramycin resistant *Streptomyces* colonies were re-isolated from the control soil wells, indicating that any resulting colonies arising from the treated wells were derived from inoculated strains. A generalised linear model (GLM) with a negative binomial distribution demonstrated that, overall, a significantly greater number of *S. coelicolor* M145-eGFP colonies could be recovered from soil wells than *Streptomyces* M3-eGFP (*P* < 0.002). However, there was no significant effect of the different soil wetting treatments (either water or salicylate) on CFU number (*P* = 0.07; Fig. 3). The interaction term between strain and wetting treatment was also insignificant (*P* = 0.20), indicating that this did not differ between the two inoculated strains. These results suggest that neither *Streptomyces* strain had a competitive advantage when greater concentrations of SA were present in soil. Taking these results together, we conclude that the observed increase in colonization by strains M145 and M3 in *A. thaliana cpr5* plants is likely due to a pleiotropic effect of plant genotype on other aspects of plant growth rather than a direct effect of the increased presence of SA in root exudates and soil.

**Figure 3.**
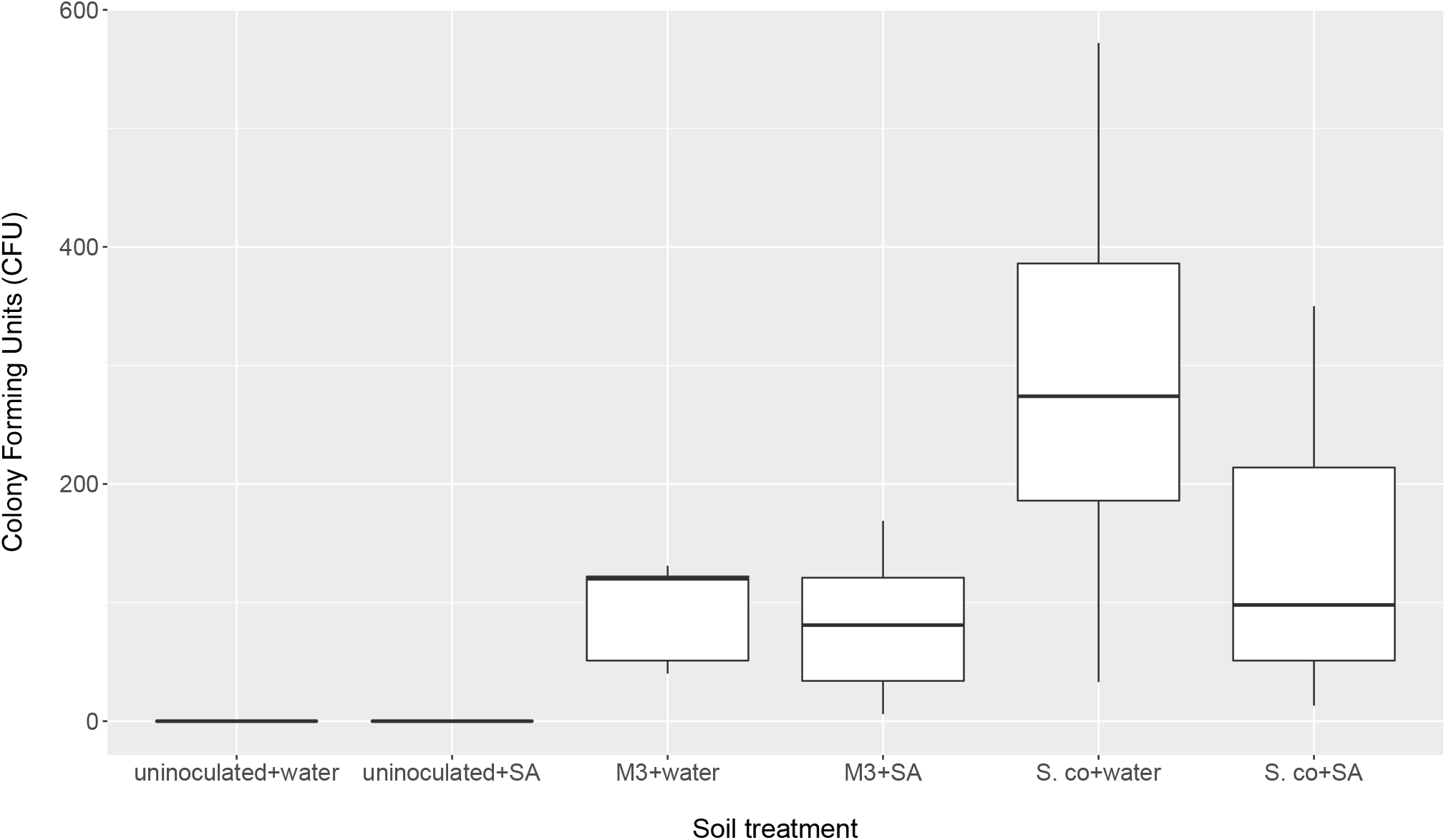
Number of colony forming units (CFU) of inoculated *Streptomyces coelicolor* M145 or *Streptomyces* M3 strains returned from soil microcosms treated with either salicylic acid (SA) or distilled water after 10 days. Control microcosms were inoculated, N = 9 microcosms per treatment.

### *Streptomyces* species feed on root exudates *in vitro* but not in soil

We then set out to test the hypothesis that *Streptomyces* bacteria can grow using a wider range of *A. thaliana* root exudate molecules as their sole nutrients source. We tested two *S. lydicus* strains, the five *Streptomyces* strains isolated from surface-washed *A. thaliana* roots (11) and the model organism *S. coelicolor* M145 (Table 1). *A. thaliana* root exudates were collected using a small-scale hydroponics system and used to make plates with purified agarose rather than agar, because streptomycetes can grow on the impurities in agar (not shown). Control plates were made using agarose and sterile water. The results show that all eight strains could grow on plates containing *A. thaliana* root exudates but not on the water-only control plates (Fig. 4) implying that all of the *Streptomyces* strains tested here can use *A. thaliana* root exudate as their sole source food source *in vitro*.

**Figure 4.**
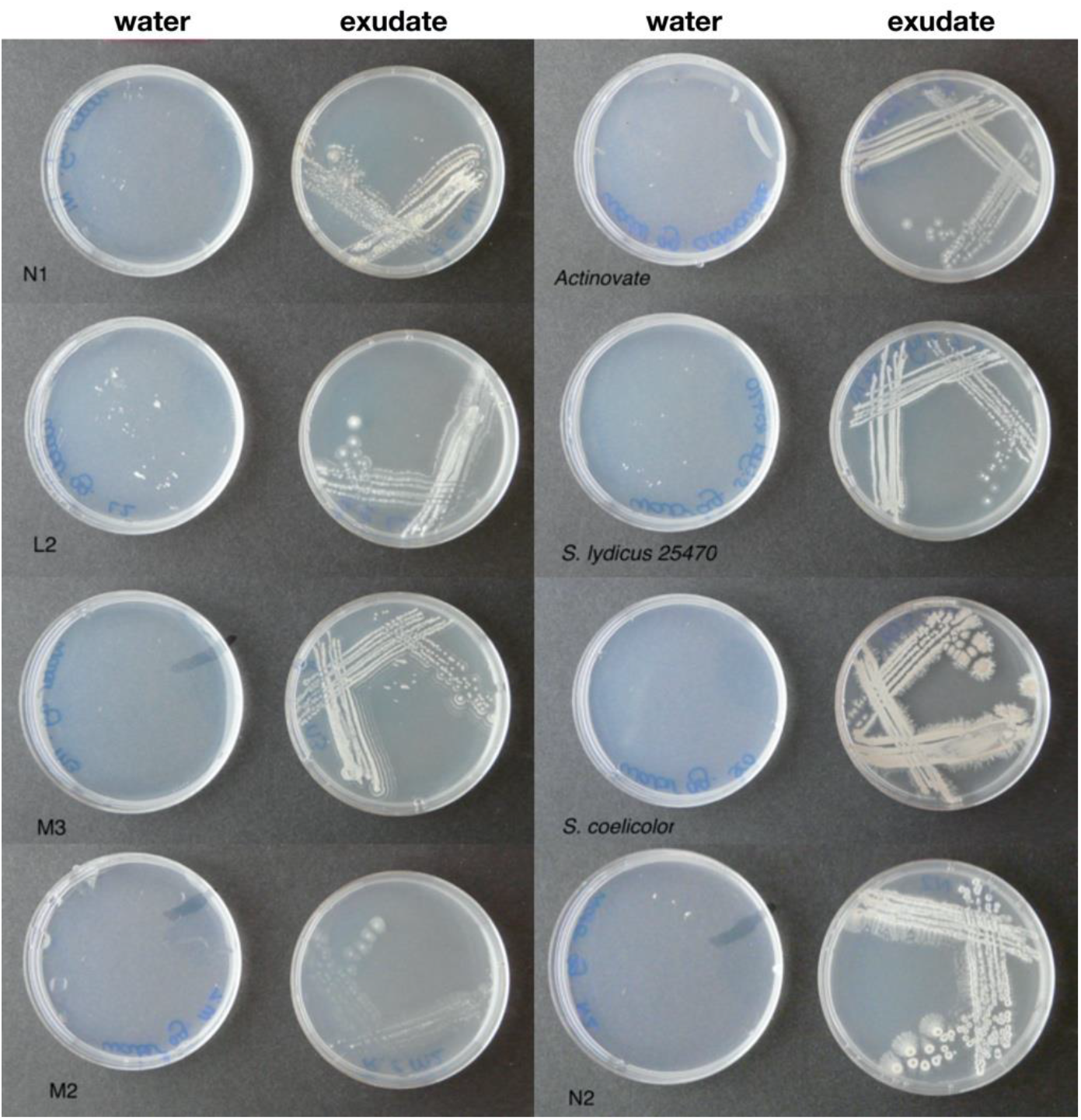
*Streptomyces* isolates growing on root exudates as a sole food source. Strains N1, L2, M3, M2 and N2 were isolated from surface washed *A. thaliana* roots. *Actinovate* is the *S. lydicus* WYEC106 cultured from the horticultural product Actinovate. *S. lydicus* 25470 was acquired from the American Type Culture Collection (see Table S1 in Additional File 1 for more details).

Several independent studies have reported that *A. thaliana* plants have a relatively stable and consistent root bacterial community, which is enriched with members of the phylum Actinobacteria; this enrichment is predominantly driven by the presence of the family *Streptomycetaceae* (4–8,10,15). To test whether this was the case for plants grown in our laboratory, we repeated the bacterial 16S rRNA gene profiling using universal primers PRK341F and PRK806R (Table S2). Plants were grown under controlled conditions in Levington F2 compost to match the conditions in our previously published study (11) and the other experiments reported here (for chemical analysis of the Levington F2 compost, see Table S3). We found that, in agreement with published studies, *Streptomycetaceae* was the most abundant family of Actinobacteria in both the rhizosphere and root compartment of *A. thaliana* plants (Fig. 5), making up 5.62% (± 1.10% standard deviation) of the rhizosphere community and 1.12% (± 0.50%) of the endophytic community, respectively. Actinobacteria as a whole made up 15.15% (± 1.47%) and 2.87% (± 0.88%) of these compartments, respectively (Figs 5 and S2). However, bacteria of the phylum Proteobacteria were found to dominate in all of the soil, rhizosphere and root compartments of the *A. thaliana* plants (Fig S2) and were also significantly enriched in the endophytic compartment, relative to the surrounding soil and rhizosphere (*P* <0.05 in Dunn’s multiple comparison tests between root and rhizosphere and root and soil compartments) increasing to 91.22% (± 0.74% standard deviation) of the root-associated community, compared to 38.85% (± 9.80%) of the rhizosphere and 35.46% (± 1.97%) of the soil community, respectively.

**Figure 5.**
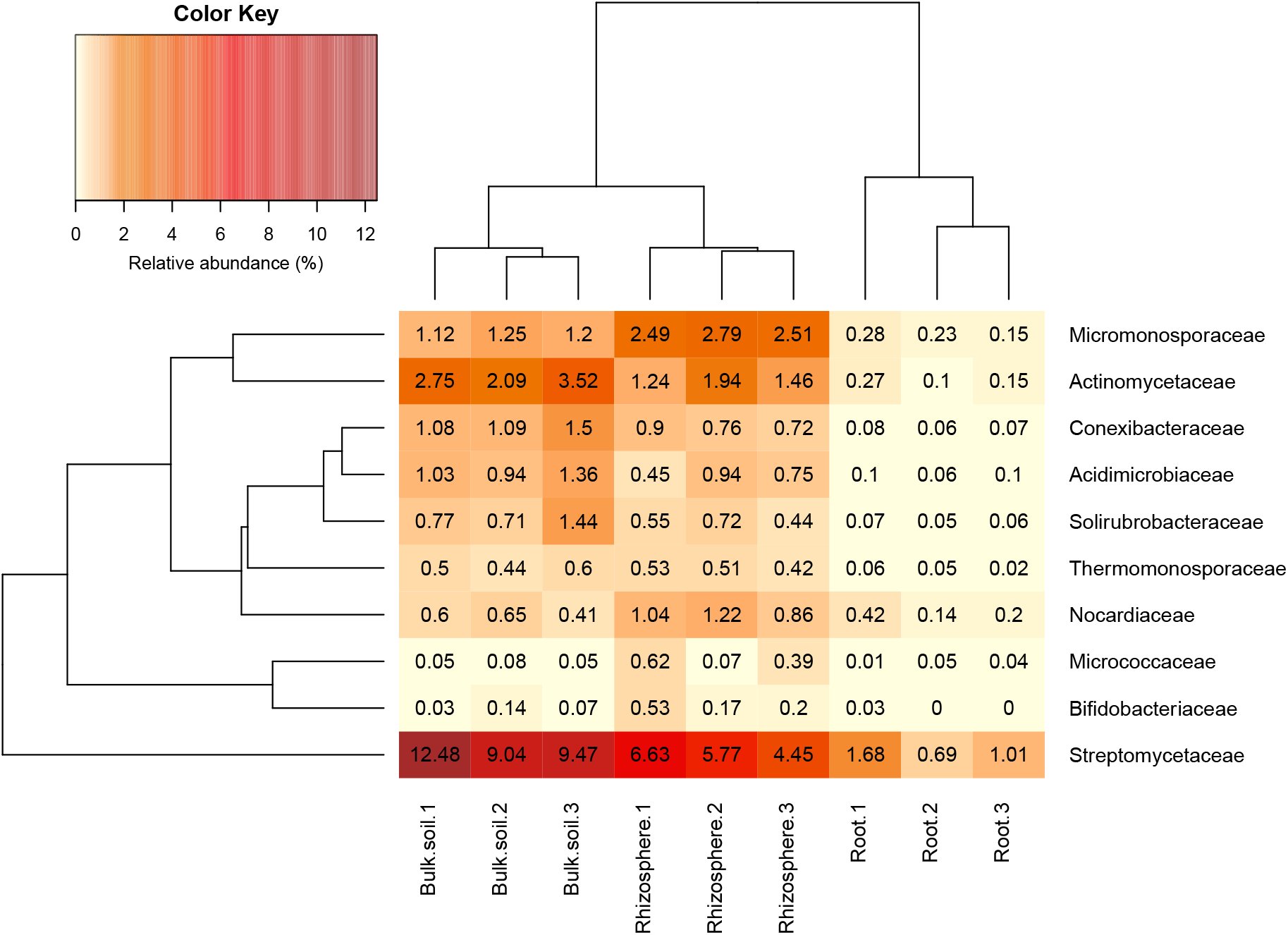
The relative abundance of actinobacterial families in soil, rhizosphere and root compartments. N = 3 replicate plants in individual pots. Streptomycetaceae was the most abundant family of Actinobacteria in all three compartments. Clustering represents Bray Curtis dissimilarities.

To test whether *Streptomyces* species metabolize *A. thaliana* root exudates we used ^13^CO_2_ DNA-SIP (16,31,32). Plants incubated with ^13^CO_2_ fix the ‘heavy’ isotope of carbon which then becomes incorporated into carbon-based metabolites; many of these then leave the plant as root exudates. Bacteria feeding off labelled root exudates in the rhizosphere or plant metabolites in the endosphere will incorporate the ^13^C into their DNA as they grow and divide. By separating ^12^C- and ^13^C-labelled DNA on a cesium chloride gradient and using universal primers (Table S2) to amplify and sequence a variable region of the 16S rRNA gene from both heavy and light fractions, we can determine which bacteria are being fed by the plant in the rhizosphere and endosphere of *A. thaliana* (16,33). *A. thaliana* plants were labelled by growing them (n=3) in Levington F2 compost for 21 days in sealed growth chambers in the presence of either ^12^CO_2_ or ^13^CO_2_ (Fig. S3). Unplanted pots were also incubated with ^13^CO_2_ to account for autotrophic metabolism. Given the relatively short time frame of CO_2_ exposure, ^13^C was expected to be predominantly incorporated into plant metabolites rather than into plant cell wall material. Bacteria that incorporate ^13^C into their DNA must therefore either be feeding on plant metabolites or fixing ^13^CO_2_ autotrophically. After 21 days, total DNA was extracted from the rhizosphere, endosphere and unplanted soil compartments of the ^12^CO_2_ or ^13^CO_2_ incubated plants and heavy (^13^C) and light (^12^C) DNA were separated by density gradient ultracentrifugation. For the root samples, it was necessary to combine the three replicate samples into one sample for each of the CO_2_ treatments before ultracentrifugation, due to low DNA yields. Heavy and light DNA fractions of each gradient were determined via qPCR and used for 16S rRNA gene amplicon sequencing (Fig. S4).

A principle coordinates analysis, using a Bray-Curtis dissimilarity matrix of the relative abundances of genera present in each of the different rhizosphere fractions suggested that certain genera had metabolised ^13^C labelled host-derived resources, since ^13^C and ^12^C heavy fractions separated spatially on an ordination plot, indicating differences between the bacterial communities in these fractions (Fig. S5). A permutational analysis of variance (PERMANOVA) analysis confirmed that fraction type had a significant effect on the community composition (permutations=999, R^2^= 0.80, *P* = <0.01). A total of 28 genera showed an average of two-fold (or more) enrichment in the ^13^C heavy (^13^CH) fraction of rhizosphere samples, compared to both the ^13^CO_2_ light fraction (^13^CL) and ^12^CO_2_ heavy fraction (^12^CH), respectively, suggesting they had become labelled through the metabolism of root exudates (Table S4; Fig. S6). Importantly, the abundance of these bacteria was not enriched in heavy versus light fractions of ^13^CO_2_ unplanted controls, meaning they were not autotrophically fixing CO_2_. The majority of these taxa were in the phylum Proteobacteria (24 out of 28 genera), with only one representative each from the phyla Chloroflexi (*Levilinea*), Firmicutes (*Pelotomaculum*), Cyanobacteria (*Chroococcidiopsis*), and the Planctomycetes (*Pirellula*) (Table S4). The most enriched genus was *Pseudomonas*, which demonstrated a 64-fold enrichment in relative abundance between the ^13^CH (8.63% ± 3.33% average relative abundance ± standard error) and ^13^CL (0.13% ± 0.01%) fractions and a 23-fold enrichment between the ^13^CH fraction compared to the ^12^CH control (0.37% ± 0.06%) (Fig. S6).

A total of 27 genera demonstrated more than a two-fold enrichment in the ^13^CH fraction of the endophytic compartment versus ^13^CL and ^12^CH fractions (Fig. 6). The majority of these genera were also Proteobacteria (21 genera in total), with three genera belonging to the phylum Planctomycetes (*Blastopirellula, Pirellula* and *Gemmata*), one to the Firmicutes (*Clostridium*), one to the Chloroflexi (*Chloroflexus*), and one to the Actinobacteria (*Jatrophihabitans*) (Table S4). The most abundant genus in the ^13^CH fraction of the endophytic samples was *Shinella*, which demonstrated a 38-fold enrichment between the ^13^CH (26.42% relative abundance) and ^13^CL (0.68% relative abundance) fractions and a 23-fold enrichment between the ^13^CH and the ^12^CH control fractions (1.14%) (Fig. 6). In terms of fold change, however, *Pseudomonas* was the most enriched genus with a 26-fold increase in abundance between ^13^CH (15.64% relative abundance) and ^13^CL (0.59% relative abundance) fractions of ^13^CO_2_ incubated plants (Fig. 6). 15 out of the 28 genera showed more than a two-fold enrichment in the heavy fraction of the endosphere but not the rhizosphere samples, whereas 16 out of 29 genera were metabolising exudates in the rhizosphere but not the endosphere. The remaining 13 genera were enriched in the heavy fractions of both the endophytic and rhizospheric compartments, suggesting that they are able to survive and make use of plant metabolites in both of these niches (Table S4).

**Figure 6.**
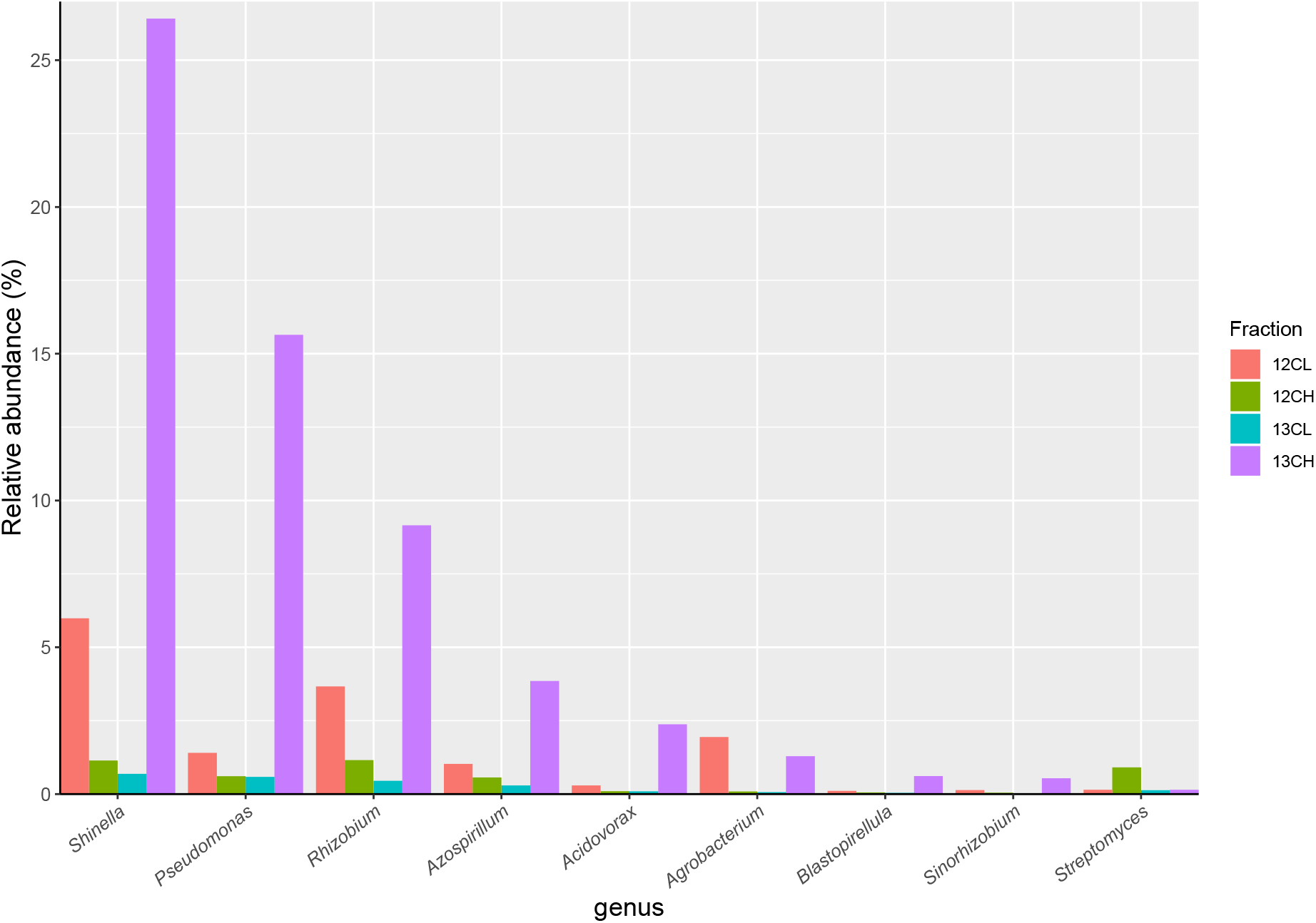
The relative abundance of eight genera of bacteria that demonstrated the greatest overall enrichment between the ^13^CH fraction and the ^12^CH fraction due to the metabolism of ^13^C labelled root exudates in the endophytic compartment of *Arabidopsis thaliana* plants. In comparison, the genus *Streptomyces* did not demonstrate enrichment between these fractions suggesting they did not metabolise ^13^C-labelled root exudates. 12CL/13CL and 12CH/13CH are the buoyant density fractions representing the light/unlabelled (L) and heavy/labelled (H) fractions of DNA from ^12^CO_2_ (12C) or ^13^CO_2_ (13C) incubated plants, respectively.

Surprisingly, there was no enrichment of *Streptomyces* bacteria in the ^13^CH fractions of either the rhizosphere or endosphere, despite being the most dominant member of the phylum Actinobacteria in both compartments (Figs 6, S6 and S7; Table S4). Taken together, these data suggest that, in compost, *Streptomyces* bacteria are outcompeted for root exudates by unicellular bacteria, particularly members of the phylum Proteobacteria. Proteobacteria were abundant in the unfractionated soil (35% average relative abundance), rhizosphere (39%) and endophytic compartment (91%), compared to Actinobacteria (present at 20%, 15% and 3% in the soil, rhizosphere and root compartments, respectively) (Fig. S2) and therefore may have the upper hand at the outset of competition [33]. Accordingly, genera that were found to be metabolizing the greatest amount of exudates in the roots were also found to be enriched in the endophytic compartment compared to the surrounding soil (Fig. S2 and S7).

## Discussion

Although typically described as free-living soil bacteria, there is increasing evidence that *Streptomyces* species are effective at colonizing the rhizosphere and endosphere of plant roots where they can promote plant growth and provide protection against disease (11–13,34,35). It has even been suggested that their diverse specialized metabolism and filamentous growth may have evolved to give them a competitive advantage in this niche since it occurred ~50 million years after plants first colonized land (3). Plants certainly provide a good source of food in nutritionally poor soil environments, both through the release of root exudates but also by shedding more complex organic cell wall material as roots grow and senesce (36). There is growing evidence that root exudates enable plants to attract bacteria and shape their microbiome (17,37–40). However, although we confirmed in this study that *Streptomyces* bacteria can survive using only *A. thaliana* root exudates *in vitro* (Fig. 4), ^13^CO_2_ DNA-SIP revealed that they were not ^13^C-labelled in our experiments and were in fact outcompeted for plant metabolites in both the rhizosphere and endosphere compartments by proteobacterial genera, that were abundant in both compartments (Figs 6 and S6). Many proteobacteria are thought to be “r-strategists” that have the ability to thrive and rapidly proliferate in fluctuating environments where resources are highly abundant, such as in the rhizosphere and root-associated microbiome (41). In contrast, species of *Streptomyce*s typically exhibit slower growth and are competitive in environments where resource availability is limited, such as in the bulk soil (42). Thus, it is possible that proteobacteria were able to rapidly proliferate around and within the *A. thaliana* plant roots and outcompete the slower growing actinobacterial species, particularly those that were already present at lower abundance. Given their diverse metabolic capabilities, it is likely that streptomycetes are able to persist at low abundance by feeding on more complex organic polymers under conditions of high competition, including older, plant-originated organic material such as root cells and mucilage that were sloughed off before ^13^C labelling commenced. *A. thaliana* also appears to exhibit a relatively small rhizosphere effect compared to other plant species (7,14), meaning that there is a weaker differentiation in terms of microenvironment and community composition between the rhizosphere and bulk soil, than there is between these two compartments and the endophytic niche, as observed in this study (Fig. S2). Thus, complex polymers may have been more readily available in the rhizosphere than exudates, particularly given the fact that the compost growth medium also contained a relatively high level of organic matter (91.08%, Table S3).

We also tested whether *Streptomyces* strains *S. coelicolor* M145 and the *A. thaliana* endophyte *Streptomyces* M3 are specifically attracted to *A. thaliana* roots by a specific root exudate compound, salicylate, since a previous study suggested this molecule can modulate microbiome composition and specifically attract streptomycetes (10). However, we found no evidence to support this hypothesis. We found that *cpr5* mutant plants, which constitutively produce and accumulate salicylate, are colonized more readily by streptomycetes than wild-type plants, but we found no difference between the levels of *Streptomyces* root colonization in wild-type *A. thaliana* and salicylate-deficient *sid2-2* and *pad4* plants. Our data suggest that the *cpr5* effect is not directly due to salicylate, at least with the M145 and M3 strains we tested here. None of the eight *Streptomyces* strains we tested could use salicylate as a sole carbon source, and they did not respond chemotactically to salicylate or compete better in salicylate amended soil. We propose that the greater colonization efficiency observed in *A. thaliana cpr5* plants in this and a previous study (10) was not due to increased levels of salicylate, but more likely due to one or more of the pleiotropic effects of the *cpr5* mutation (21). The *cpr5* gene encodes a putative transmembrane protein (28) that is part of a complex regulatory signaling network; its deletion results not only in the constitutive activation of SA biosynthesis genes, but also stunted root and aerial growth, disrupted cellular organization, spontaneous cell death, early senescence in young plants, lesions on various tissue types, disrupted cell wall biogenesis, an increased production of ROS and high levels of oxidative stress (21,26–28,43,44). Thus, it is possible that the altered cell wall composition and spontaneous cell death in *crp5* plants could have resulted in easier access to plant roots by streptomycetes, particularly as these bacteria are thought to enter roots in compromised areas, such as lesions and sites of wounding (12,45). We also note that although that original study (10) reported that salicylate could be used as a sole C and N source by a *Streptomyces* endophyte strain, their supplementary information reported that sodium citrate was included in the growth medium used in these experiments and streptomycetes can grow using sodium citrate alone (Fig. 2).

In summary, we conclude that *Streptomyces* bacteria do not appear to be attracted to the rhizosphere by salicylic acid and did not feed on root exudates under the conditions presented in this study, because they were outcompeted by faster growing proteobacteria. Instead, it is likely that streptomycete bacteria were able to persist in the root microbiome by utilising the organic matter present in the soil growth medium in addition to components of the plant cells (such as cellulose) that are sloughed off during root growth. These ubiquitous soil bacteria are likely to be present in the soil as spores or mycelia when plant seeds germinate and root growth is initiated, enabling successful transmission between plant hosts. Although not tested in this study, it would be interesting to see whether *Streptomyces* spores germinate in the presence of plant root exudates. Understanding the factors that attract beneficial bacteria to the plant root microbiome, and maintain them there, may have significant implications for our ability to develop and deliver effective plant growth-promoting agents that are competitive within the plant root niche. Indeed, a range of studies have shown that organic matter amendments can improve the competitiveness and overall effectiveness of streptomycete biocontrol strains, presumably by providing them with an additional source of nutrients (13). Clearly some *Streptomyces* strains can colonize the endosphere of plants and confer advantages to the plant host but it is not yet clear what advantage these bacteria gain from entering the plant roots, given that they must have to evade or suppress the plant immune system and are outcompeted for plant metabolites inside or outside the roots. Future work will address these questions and test the hypothesis that soil is a mechanism for vertical transmission of *Streptomyces* between generations of plants.

## Materials and Methods

### Isolation and maintenance of *Streptomyces* strains

The *Streptomyces* strains L2, M2, M3, N1 and N2 (Table 1) were isolated from surface-washed *A. thaliana* roots in a previous study (11). Genome sequences were also generated for each of these isolates, as well as for three strains of *Streptomyces lydicus*, one of which was isolated from the commercial biocontrol product Actinovate, whilst the remaining two (ATCC25470 and ATCC31975) were obtained from the American Type Culture Collection (Table 1). In the present study, *Streptomyces* strains were maintained on SFM agar (N1, N2, M2, M3 and *S. coelicolor* M145), Maltose/Yeast extract/Malt extract (MYM) agar with trace elements (L2) or ISP2 agar (*S. lydicus* strains) (Table S5). Strains were spore stocked as described previously (46).

### Generating eGFP-labelled *Streptomyces* strains

Plasmid pIJ8660 (Table S1) containing an optimized eGFP gene and an aac apramycin resistance marker (47) was altered using molecular cloning of the constitutive *ermE\** promotor to drive eGFP expression. Primers ermEKpnFwd and ermENdeRev (Table S1) were used to amplify *ermEp\** from the plasmid pIJ10257 (Table S1)(48). The PCR product was subsequently purified and then digested with the restriction enzymes KpnI and NdeI, before being ligated into the same sites in pIJ8660. The resulting plasmid, called pIJ8660/*ermEp\** (Table S1), was conjugated into the *Streptomyces* strains M3 and *S. coelicolor* M145 (Table 1) as described previously (46). Exconjugants were selected and maintained on SFM agar plates containing 50 mg ml^−1^apramycin.

### Plant root colonization assays in soil

Wild-type *A. thaliana* Col-0 as well as the mutant plant lines *cpr5, pad4*, and *sid2-2* were obtained from the Nottingham *Arabidopsis* Stock Centre (Table 1). Seeds were sterilized by placing them in 70% (v/v) ethanol for 2 minutes, followed by 20% NaOCl for 2 minutes and then washing them five times in sterile dH_2_O. Spores of either *S. coelicolor* M145-eGFP or *Streptomyces* M3-eGFP (Table 1) were used as inoculants and were pre-germinated in 2xYT (Table S5) at 50°C for 10 minutes (46). Uninoculated 2xYT was used as a control. Surface sterilized seeds were placed into 500 μl 2xYT containing 10^6^ spores ml^−1^ of pre-germinated *Streptomyces* spores; these were mixed for 90 minutes on a rotating shaker before being transferred to pots of sieved Levington F2 seed and modular compost, soaked with distilled H_2_O. Another 1 ml of pre germinated spores (or 2xYT for control pots) was pipetted into the soil to a depth of approximately 2 cm below the seed. Pots were incubated at 4°C for 24 hours, then grown for 4 weeks under a 12-hour photoperiod. Six replicate plants were grown for each plant genotype and each *Streptomyces* strain.

### Re-isolation of *Streptomyces* bacteria from roots

In order to re-isolate tagged strains from the roots of the different *A. thaliana* genotypes we used a previously established method (10). Briefly, plants were taken aseptically from pots and their roots were tapped firmly to remove as much adhering soil as possible. Root material was washed twice in sterile PBS-S buffer (Table S5) for 30 minutes on a shaking platform and then any remaining soil particles were removed with sterile tweezers. Cleaned roots were then transferred to 25ml of fresh PBS-S and sonicated for 20 minutes to remove any residual material still attached to the root surface; this ensured that any remaining bacteria were either present in the endophytic compartment or were very firmly attached to the root surface (“the rhizoplane”). Cleaned roots were weighed, then crushed in 1 ml of sterile 10% (v/v) glycerol. 100 μl of the homogenate was plated onto three replicate soya flour plus mannitol agar plates containing 50 μg ml^−1^ apramycin to select for inoculated streptomycetes (Table S5). Plates also contained 5 μg ml^−1^ nystatin and 100 μg ml^−1^ cyclohexamide to inhibit fungal growth. Agar plates were incubated at 30°C for 5 days, before the number of apramycin-resistant streptomycete colony forming units (cfu) were counted. This method has been used previously in other studies (24,25) to estimate the root colonization dynamics of streptomycetes. Counts were converted to cfu per gram of root tissue and log-transformed to normalize residuals. Data were analysed via ANOVA and post-hoc Tukey’s Honest Significant Difference (HSD) tests.

### Testing for sole use of carbon and nitrogen sources

The *A. thaliana* endophyte *Streptomyces* strains L2, M2, M3, N1 and N2, as well as the *S. lydicus* strains ATCC25470, ATCC31975 and Actinovate (Table 1), were streaked onto minimal agarose medium plates supplemented with either their preferred carbon source as a positive control, 3.875 mM sodium citrate, or 0.5 mM salicylic acid (SA) as a carbon source (Table S5). Preferred carbon sources were 5 g L^−1^ of mannitol for strains N2 and M2; 5 g L^−1^ maltose for strains M3, N1, *S. lydicus* 25470 and *S. lydicus* Actinovate; 5 g L^−1^ sucrose for L2; 10 g L^−1^ glucose for *S. lydicus* 31975. Plates with no carbon source were used as a negative control. All strains were plated in triplicate onto each type of media and incubated for seven days at 30°C before imaging. *S. coelicolor* M145 was not tested as it can use agarose as a sole carbon source (49,50). Agarose was used as a gelling agent as streptomycetes can grow on impurities that are present in agar.

### Screening for salicylate metabolism genes in the genomes of *Streptomyces* isolates

To identify whether the nine different streptomycetes carried homologues to known SA degradation genes, all experimentally verified pathways involving SA (or salicylate) degradation were identified using the MetaCyc database (http://www.metacyc.org/) and amino acid sequences of characterized genes involved in each of the five pathways were then retrieved from UniProt (Table S1). Protein sequences were used to perform BLASTP searches against predicted Open Reading Frames (ORFs) for each genome-sequenced streptomycete. Previously generated sequences for N1, N2, M2, M3, L2, *S. lydicus* 25470, *S. lydicus* 31975 and *S. lydicus* Actinovate were used (11) as well as the published genome sequence of *S. coelicolor* (51). The results of the best hit (% identity and % query coverage) are reported.

### SA as a chemoattractant

To test whether streptomycetes grew towards SA, 4 μl of spores (10^6^ spores ml^−1^) of each streptomycete (Table 1) were pipetted onto the centre of SFM agar plates. 40 μl of either 1 mM or 0.5 mM filter-sterilized SA was inoculated onto 6 mm filter paper discs (Whatman) and allowed to dry. Discs were then added to one side of the agar plate, 2 cm from the streptomycete spores. SA solutions were prepared by diluting a 100 mM stock solution to a 1 mM or 0.5 mM SA solution in PBS. The 100 mM stock solution was made by dissolving 0.138 g of SA in 2 ml of 100% DMSO before making the solution up to 10 ml with PBS. Thus, the resulting 1 mM and 0.5 mM solutions had a final concentration of 0.2% (v/v) and 0.1% (v/v) DMSO, respectively. To check that any observations were not due to the effects of DMSO, control plates were also run alongside the SA experiment, in which discs were soaked in 40 μl of a 0.2% DMSO (in PBS), equivalent to the final concentration of DMSO in the 1 mM SA solution. All strains and disc type pairings were plated in triplicate and were incubated for seven days at 30°C before imaging.

### Enumeration of bacteria from soil microcosms following exogenous application of SA

Levington F2 compost (4 ml) was placed into each compartment of a 12-well plate and soaked with 0.5 ml of sterile dH_2_O or 0.5 mM SA. Each well was then inoculated with a spore solution (10^7^ spores ml^−1^) of either *S. coelicolor* M145-eGFP or *Streptomyces* M3-eGFP (Table 1), suspended in dH_2_O or 0.5 mM SA. Spores of eGFP-tagged M3 or M145 were used to align with *in vivo* plant colonization experiments (outlined above). There were nine replicate wells for each treatment. Well-plates were placed under a 12-hour photoperiod for 10 days. 100 mg of soil from each well was then diluted in 900 μl of water and vortexed. Serial dilutions were then plated onto SFM containing 50 μg ml^−1^ apramycin (for selection) in addition to 10 μg ml^−1^ nystatin and 100 μg ml^−1^ cyclohexamide (to repress fungal growth). CFU of the *Streptomyces* inoculum were then enumerated on 10^−2^ dilution plates after 4 days to assess whether SA affected the competitiveness of strains in soil. A generalized linear model (GLM) with a negative binomial distribution was generated, using the package MASS in R 3.2.3 (52), to model the effect of strain (*S. coelicolor* M145-eGFP or *Streptomyces* M3-eGFP), soil treatment (wetting with SA or dH2O), and their interaction term (soil treatment) on the number of bacterial cfu returned from soil wells. Likelihood ratio tests were used to establish the significance of terms in the model.

### Root exudate collection and growth of Streptomycetes

To test whether streptomycetes were able to utilise other compounds in root exudates more generally, root exudates were isolated from *A. thaliana* and used as a growth medium for the *Streptomyces* strains (Table 1) that we previously isolated from *A. thaliana* plants growing in a compost system (11). Seeds of *A. thaliana* Col-0 were sterilised (as described above) and germinated on MSk agar (Table S5). After 7 days of growth at 22°C under a photoperiod of 12 hours light/12 hours dark, seedlings were transferred to 12 well plates, containing 3 ml of liquid MSk (Table S5, 0% w/v sucrose) in each well. Seedlings were then grown for a further 10 days before being washed and transferred to new wells containing 3 ml of sterile water. After 5 days plant material was removed and the liquid from each well was filter-sterilised and added to sterile agarose (0.8% w/v) to make solid growth medium plates. Spores of previously isolated *Streptomyces* isolates (Table 1) were streaked onto these plates and incubated for 7 days at 30°C. Agarose/water (0.8% w/v) plates were used as a control.

### ^13^CO_2_ Stable Isotope Probing

Seeds of *A. thaliana* Col-0 (Table 1) were sterilised, as described above, before being sown singly into pots containing 100 ml sieved Levington F2 compost, soaked with dH_2_O. These were placed in the dark at 4°C for 48 hours, after which they were transferred to short-day growth conditions (8 h light/16 h dark) at 22°C for 32 days before exposure to CO_2_ treatments. Each plant was placed into an air-tight, transparent 1.9 L cylindrical tube. Three plants were exposed to 1,000 ppmv of ^12^CO_2_ and three plants were exposed to 1,000 ppmv of ^13^CO_2_ (99%, Cambridge isotopes, Massachusetts, USA). Three unplanted controls containing only Levington F2 compost were additionally exposed to 1000 ppmv of ^13^C labelled CO_2_ to control for autotrophic consumption of CO_2_ by soil microbes. CO_2_ treatments took place over a period of 21 days. CO_2_ was manually injected into tubes every 20 minutes over the course of the 8 hour light period, to maintain the CO_2_ concentration at ~1,000 ppmv; CO_2_ concentration was measured using gas chromatography (see below). At the end of the light period each day, tube lids were removed to prevent the build-up of respiratory CO_2_. Just before the next light period, tubes were flushed with an 80%/20% (v/v) nitrogen-oxygen mix to remove any residual CO_2_ before replacing the lids and beginning the first injection of CO_2_.

### Gas Chromatography

The volume of CO_2_ to be added at each injection in the SIP experiment was determined by measuring the rate of uptake of CO_2_ over 20 minutes every 4 days. This was done using an Agilent 7890A gas chromatography instrument with a flame ionisation detector and a Poropak Q (6ft x 1/8”) HP plot/Q (30 m x 0.530 mm, 40 μm film) column with a nickel catalyst and a nitrogen carrier gas. The instrument was run with the following settings: injector temperature 250°C, detector temperature 300°C, column temperature 115°C and oven temperature 50°C. The injection volume was 100 μl and the run time was 5 mins, with CO_2_ having a retention time of 3.4 mins. Peak areas were compared to a standard curve (standards of known CO_2_ concentration were prepared in 120 ml serum vials that had been flushed with 80%/20% nitrogen-oxygen mixture).

### Sampling and DNA extraction from soil, rhizosphere, and roots

Two samples of root-free “bulk soil” were collected from each planted and unplanted pot; samples were snap-frozen in liquid nitrogen and stored at −80°C. Root were then processed to collect the rhizosphere soil and endophytic samples according to a published protocol (4). For the planted pots, roots were tapped until only the soil firmly adhering to the root surface remained; the remaining soil was defined as the rhizosphere fraction. To collect this, roots were placed in 25 ml sterile, PBS-S buffer (Table S5) and washed on a shaking platform at top speed for 30 min before being transferred to fresh PBS-S. Used PBS-S from the first washing stage was centrifuged at 1,500 x g for 15 minutes, and the supernatant was removed. The resulting pellet (the rhizosphere sample) was snap-frozen and stored at −80°C. The roots were then shaken in fresh PBS-S for a further 30 minutes before removing any remaining soil particles with sterile tweezers. Finally, the cleaned roots were transferred to fresh PBS-S and sonicated for 20 minutes in a sonicating water bath (4). Root samples for each plant were then snap frozen and stored at −80°C. Each root sample consisted of bacteria within the roots and those very firmly attached to the root surface (the “rhizoplane”). A modified version of the manufacturer’s protocol for the FastDNA™ SPIN Kit for Soil (MP Biomedicals) was used to extract DNA from soil, rhizosphere, and root samples. Modifications included pre-homogenisation of the root material by grinding in liquid nitrogen before adding lysis buffer, an extended incubation time (10 minutes) in DNA matrix buffer, and elution in 150 μl of sterile water. DNA yields were quantified using a Qubit™ fluorimeter.

### Density gradient ultracentrifugation and fractionation

DNA samples from the rhizosphere, roots, and unplanted soil were subjected to caesium chloride (CsCl) density gradient separation using an established protocol (33). For each of the replicate rhizosphere samples from both the ^12^CO_2_ and ^13^CO_2_ incubated plants, 1.5 μg of DNA was loaded into the CsCl solution with gradient buffer. For the three unplanted soil sample replicates, 1 μg of DNA was used. For the root samples it was necessary to combine the three replicate samples for each CO_2_ treatment due to low DNA yields. Thus, 0.2 μg of DNA was pooled from each of the three replicates per ^12^CO_2_ and ^13^CO_2_ treatment, and the final 0.6 μg was loaded into the CsCl solution. After ultracentrifugation, the density of each fraction was measured using a refractometer (Reichert Analytical Instruments, NY, USA) in order to check for successful gradient formation. DNA was precipitated from fractions (33) and stored at −20°C before use as a template in qPCR and PCR reactions.

### Fraction selection via qPCR and 16S rRNA gene amplicon sequencing

To identify fractions containing heavy (^13^C) and light (^12^C) DNA for each sample, 16S rRNA gene copy number was quantified across fractions using qPCR. Reactions were carried out in 25 μl volumes. 1 μl of template DNA (either sample DNA or standard DNA), or dH_2_O as a control, was added to 24 μl of reaction mix containing 12.5 μl of 2x Sybr Green Jumpstart Taq Ready-mix (Sigma Aldrich), 0.125 μl of each of the primers PRK341F and MPRK806R (Table S2), 4 μl of 25 mM MgCl2, 0.25 μl of 20 μg μl^−1^ Bovine Serum Albumin (BSA, Roche), and 7 μl dH_2_O. Sample DNA, standards (a dilution series of the target 16S rRNA gene at known molecular quantities), and negative controls were quantified in duplicate. Reactions were run under the following conditions: 96°C for 10 mins; 40 cycles of 96°C for 30 sec, 52°C for 30 sec, and 72°C for 1 min; 96°C for 15 sec; 100 cycles at 75°C-95°C for 10 secs, ramping 0.2°C per cycle. Reactions were performed in 96-well plates (Bio-Rad). The threshold cycle (CT) for each sample was then converted to target molecule number by comparing to CT values of a dilution series of target DNA standards. Fractions spanning the peaks in 16S rRNA gene copy number were identified and equal quantities of these were combined to create a “heavy” buoyant density (labelled) and “light” buoyant density (unlabelled) fraction for each sample, respectively. The 16S rRNA genes were amplified for each of these fractions using the Universal primers PRK341F and MPRK806R (Table S2), and the resulting PCR product was purified and submitted for 16S rRNA gene amplicon sequencing using an Illumina MiSeq at MR DNA (Molecular Research LP), Shallowater, Texas, USA. Sequence data were then processed at MR DNA using their custom pipeline (53,54). As part of this pipeline, pair-end sequences were merged, barcodes were trimmed, and sequences of less than 150 bp and/or with ambiguous base calls were removed. The resulting sequences were denoised, and OTUs were assigned by clustering at 97% similarity. Chimeras were removed, and OTUs were assigned taxonomies using *BlastN* against a curated database from GreenGenes, RDPII, and NCBI (55). Plastid-like sequences were removed from the analysis. All data received from MR DNA were then further processed and statistically analysed using R 3.2.3 (52), using the packages tidyr and reshape (for manipulating data-frames), ggplot2 and gplots (for plotting graphs and heatmaps, respectively), vegan (for calculating Bray-Curtis dissimilarities, conducting principle coordinate and PERMANOVA analyses, and for generating heatmaps) and ellipse (for plotting principle coordinate analyses). All the 16S rRNA gene amplicon sequences have been submitted to the European Nucleotide Archive (ENA) database under the study accession number PRJEB30923.

### Chemical analysis of Levington compost

Soil pH, organic matter content (%) as well as the levels of phosphorous, potassium and magnesium (all in mg/kg), was measured by the James Hutton Institute Soil Analysis Service (Aberdeen, UK). To quantify inorganic nitrate and ammonium concentrations a KCl extraction was performed in triplicate, whereby 3 g of soil was suspended in 24 ml of 1 M KCl and incubated for 30 minutes, with shaking, at 250 rpm. To quantify ammonium concentration (g/kg) an indophenol blue method was used as described in (55). For nitrate concentration (in g/kg) vanadium (III) chloride reduction followed by chemiluminescence was used as described in (56).

### Data availability

All of the 16S rRNA gene amplicon sequences have been submitted to the European Nucleotide Archive (ENA) database under the study accession number PRJEB30923. Genome accession numbers are listed in Table 1.

